# GOLabeler: Improving Sequence-based Large-scale Protein Function Prediction by Learning to Rank

**DOI:** 10.1101/145763

**Authors:** Ronghui You, Zihan Zhang, Yi Xiong, Fengzhu Sun, Hiroshi Mamitsuka, Shangfeng Zhu

## Abstract

**Motivation**: Gene Ontology (GO) has been widely used to annotate functions of proteins and understand their biological roles. Currently only ¡1% of more than 70 million proteins in UniProtKB have experimental GO annotations, implying the strong necessity of automated function prediction (AFP) of proteins, where AFP is a hard multi-label classification problem due to one protein with a diverse number of GO terms. Most of these proteins have only sequences as input information, indicating the importance of sequence-based AFP (SAFP: sequences are the only input). Furthermore, homology-based SAFP tools are competitive in AFP competitions, while they do not necessarily work well for so-called *difficult* proteins, which have ¡60% sequence identity to proteins with annotations already. Thus, the vital and challenging problem now is to develop a method for SAFP, particularly for difficult proteins.

**Methods**: The key of this method is to extract not only homology information but also diverse, deep-rooted information/evidence from sequence inputs and integrate them into a predictor in an efficient and also effective manner. We propose GOLabeler, which integrates five component classifiers, trained from different features, including GO term frequency, sequence alignment, amino acid trigram, domains and motifs, and biophysical properties, etc., in the framework of learning to rank (LTR), a new paradigm of machine learning, especially powerful for multi-label classification.

**Results**: The empirical results obtained by examining GOLabeler extensively and thoroughly by using large-scale datasets revealed numerous favorable aspects of GOLabeler, including significant performance advantage over state-of-the-art AFP methods.

## 1 Introduction

The Gene Ontology (GO) was originally launched in 1998 for the consistent descriptions of gene and gene product (such as protein and RNA) across all species [2]. Currently GO has more than 40,000 biological concepts over three domains: Molecular Function Ontology (MFO), Biological Process Ontology (BPO) and Cellular Component Ontology (CCO). Annotating protein function by GO is crucial and useful for understanding the nature of biology. With the development of next generation sequencing technology, we have seen the explosive increase of protein sequences, while the number of proteins with experimental GO annotations is limited, due to the high time and financial cost of biochemical experiments. In fact, only less than 1% of 70 million protein sequences in UniProtKB [29] have experimental GO annotations. To reduce this huge gap, an imperative issue would be efficient automated function prediction (AFP) [25, 14].

AFP is a large-scale multi-label classification problem [33] by regarding one GO term as a class label and also one protein as an instance with multiple labels (multiple GO terms). AFP is very challenging due to: 1) structured ontology: GO terms are node labels of a directed acyclic graph (DAG), by which one gene annotated at one node must be labeled by GO terms of all ancestor nodes in the DAG; 2) many labels per protein: we checked GO terms in Swissprot [3] (Oct. 2016) with 66,841 proteins and found that a human protein is labeled by around 71 GO terms on average; 3) large variation in the number of GO terms per protein: also we found that only 634, out of all 10,236 GO terms in MFO (GO Ontology in Jun 2016), are associated with more than 50 proteins. This means that most of GO terms are associated with only a small number of proteins.

To advance the development of AFP, the first and second Critical Assessment of Functional Annotation (CAFA1 and CAFA2) challenges^1^ (competitions) were held in 2010-2011 and 2013-2014, respectively [25, 14]. After query proteins are given initially, CAFA has the deadline of submitting prediction. For a domain (MFO, BPO or CCO) in question, CAFA focuses on query proteins that have no experimental annotations before the submission deadline. A target protein is called a *limited-knowledge* protein, if it has experimental annotation in another domain; otherwise, it is called a *no-knowledge* protein [14]. Importantly, this means *no-knowledge* proteins have only sequences and no experimental annotations for all domains before the deadline. Obviously in practice majority (around 99% or more) of proteins have only sequences, meaning that AFP for *no-knowledge* proteins would be more important. In fact, experimental information, such as protein-protein interactions, are more costly than sequencing, resulting in literally limited knowledge (vastly missing information) among the large-scale data of AFP. Also sequences are less noisy than experimental data such as gene expression and can be the primary data available for various species. These would be also the reason why sequence-based AFP is important and should be tackled first.

The results of CAFA for no-knowledge benchmark have shown that simple homology-based methods with BLAST and PSI-BLAST are very competitive [1, 9, 11]. This indicates that sequence identity is even at the present time still a key to achieve high performance of sequence-based AFP, and also implies that prediction would be hard for sequences of low identities with any other sequences. For example, the *no-knowledge* benchmark can be divided into two types, according to the largest *global sequence identity* of the corresponding sequence to any other sequences in the training data [14]. That is, one type, called *difficult*, includes those with the largest sequence identity of less than 60%; otherwise they are called *easy.* So an urgent issue for AFP is, instead of relying on sequence homology only, to develop an approach which can predict the function of the *difficult* type of proteins within the scope of *sequence-based* approach. An important point of this approach would be to collect not only homology-related information but also various types of information available from sequences as diverse as possible and develop a method which can integrate all of these information effectively and also efficiently.

We propose a new method which we call *GOLabeler* for predicting functions of *no-knowledge* proteins, particularly for those in the *difficult* type. The basic idea of GOLabeler is to integrate different types of sequence-based evidence in the framework of “learning to rank” (LTR) [17]. LTR was originally developed in information retrieval (IR) for ranking web pages to be consistent with the relevance between the web pages and user queries. More generally, LTR is useful for multi-label classification, where multiple labels can be assigned to one instance, and LTR solves this problem by ranking labels and choosing top of them. Ranking web pages in IR requires a ranked outcome definitely, which is exactly the output of LTR. In AFP, currently GO terms for one protein are not ranked, while they can be ranked by some nature behind GO terms, such as if they are major or minor, and/or the relevance to the corresponding protein. So for AFP, LTR is already currently useful as a tool for multi-label classification and also very promising for the future AFP. Another noteworthy advantage of LTR is that GOLabeler can integrate multiple sequence-based evidence effectively, which are generated by different types of classifiers (or components), where all information are derived from the sequences only. This feature makes GOLabeler predict the functions of proteins having sequences with global sequences identity of less than 60% *(difficult* proteins) better than homology-based methods reasonably.

We examined the performance of GOLabeler extensively by using large-scale datasets, which were generated by approximately following the procedure of CAFA. Particularly, we compared the performance of GOLabeler with all component methods, three ensemble approaches and two sequence-based methods. The experimental results indicate significant performance advantage of GOLabeler over all competing methods in all experimental settings. In particular, the advantage of GOLabeler could be seen in the prediction for *difficult* proteins. Finally, we present a typical result by competing methods, from which we could see that GOLabeler could predict the largest number of all GO terms correctly as well as correctly predict the deepest GO terms in DAG of GO, among all competing methods.

## 2 Related work

A lot of biological information sources, such as protein structures, protein-protein interactions [19] and gene expression [31], are useful for AFP, while majority of proteins have no such information except sequences [26]. We thus focus on sequence-based approaches, in which sequences or their parts are used in various ways, such as 1) sequence alignment, 2) domains and motifs, and 3) features, etc., as follows: 1) sequence alignment: BLAST and/or PSI-BLAST are used to find homologous sequences and transfer their functional annotations to the query protein. For example, GoFDR [10], a top method in CAFA2, uses multiple sequence alignment (MSA) to generate position-specific scoring matrix (PSSM) for each GO term, to score the query against the corresponding GO term. 2) domains and motifs: they are usually functional sites of a query protein, and all of them in the query sequence are detected by using protein domain/motif resources, such as CATH [7], SCOP [22] and Pfam [28], to understand the function of the query protein. 3) features: the protein query is an amino acid sequence, from which biophysical and biochemical attributes can be generated, which can be closely related with domains, motifs and protein families but not necessarily the same as them. ProFET (Protein Feature Engineering Toolkit) [23] is a typical tool for extracting hundreds of such sequence-derived features including elementary biophysical ones. We note that these various sequence-based approaches play different roles, being complement to each other for AFP.

Thus, integrating different types of information or classifiers trained from them would be a key to improve the performance of AFP. In fact in AFP, several approaches of using the idea of integrating data/classifiers have already been proposed. MS-kNN, a top method in CAFA1 and CAFA2, predicts the function by averaging over the prediction scores from three data sources: sequences, expression and protein-protein interaction [16] (Note that MS-kNN is NOT a sequence-based method). Also Jones-UCL, the top team of CAFA1, integrates prediction scores from multiple methods by using the ‘consensus’ function (given in Eq. (1)) [6]. Recently, different data integration methods, mainly ‘one vote’, ‘weighted voting’ and ‘consensus’ (where ‘one vote’ relies on the classifier with the maximum confidence only while ‘weighted voting’ weights over input classifiers and examples are [16] and [15]), were compared [30]. One interesting finding of this work was, because of extensive complementarity among classifiers, ‘one vote’ performed very well in cross validation under their settings, which were totally different from CAFA. It would be interesting to check if this observation holds under practical large-scale data with diverse species in the CAFA setting.

All such data integration methods are not based on any advanced machine learning ideas, and this might underestimate the performance of integration. Our proposed approach, GOLabeler, is based on Learning to Rank (LTR), a rather new paradigm in machine learning, to integrate multiple classifiers trained from different sequence-derived data. Recently LTR has been effectively used in a wide variety of applications, not only information retrieval (IR), from which LTR was developed, but also particularly those in bioinformatics, such as annotating biomedical documents [18, 24] and predicting drug-target interactions [32]. LTR integrates the prediction results of component classifiers so that GO terms more relevant to the query protein should be ranked higher. One advantage of LTR is that the prediction results of components can be simply encoded as the input features of the model, over which any cutting edge classification/regression algorithm can be run. Also ranking would be somewhat close to the idea of ‘one vote’ (or top multiple votes), implying suitability to excessive complementarity of diverse sequence-derived information, while integration of classifiers is still an important feature of LTR. Thus, GOLabeler provides a nice framework of integrating different sequence-based information of AFP, and also the high possibility of improving the performance of the current AFP of *no-knowledge* proteins.

## 3 Methods

### 3.1 Notation

Let *D* be the given training data with *N_D_* proteins, i.e. |*D*| = *N*_D_. Let *G* _*i*_ be the *i*-th GO term, and *N*_*Gi*_ be the number of proteins with *Gi* in D (Note that this number is obtained by considering the structure of GO. That is, if *G*_*i*_ is assigned to a protein, this protein is with all GO terms of the ancestors of *G*_*i*_ in GO.). Let *T* be the given test data (the number of proteins: *Nt* = |*T*|), in which let *Pj* be the j-th protein. Let I(*G*_*i*_, *p*) be a binary indicator, showing if protein *p* is with ground-truth (true) *G*_*i*_. That is, if *p* has ground-truth (true) *G*_*i*_, *I* (*G*_*i*_, *p*) is one; otherwise zero. Let *S* (*G*_*i*_, *Pj*) be the score (obtained by a method), showing that *Pj* is with G_*i*_. In particular, in ensemble methods, *S*_*k*_ (*G*_*i*_, *Pj*) is the predicted score between G _*i*_ and *P* _*j*_ by the *k*-th method (component).

### 3.2 Overview

Fig 1 shows the entire scheme of GOLabeler for AFP. In testing, given the sequence of a query protein, candidate GO terms are generated from five components, which are already trained by using different types of information. Each candidate GO term receives prediction scores from the five components, resulting in a feature vector of length five. Then candidate GO terms, i.e. feature vectors, are put into the learning to rank (LTR) model, which is also already trained by using training data, and finally, a ranked list of GO terms is returned as the final output of GOLabeler.

**Figure 1:**
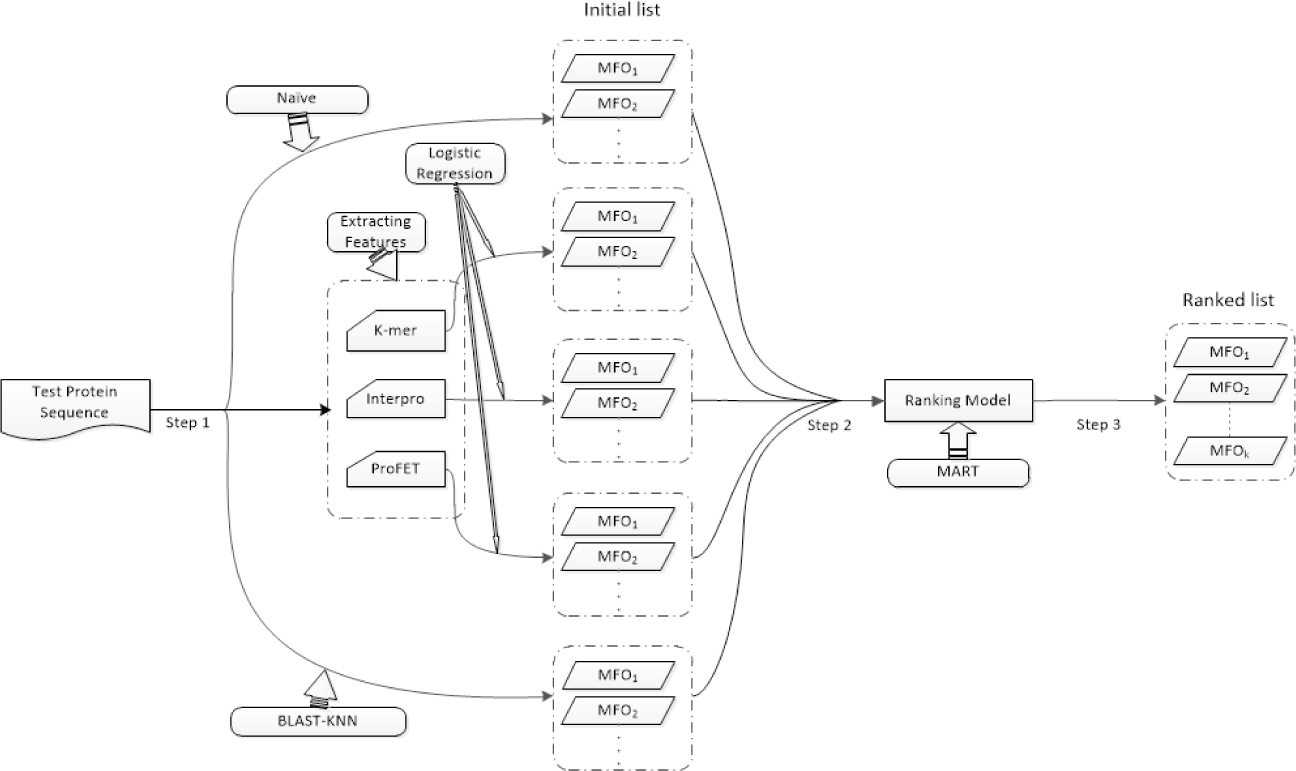
Entire scheme of GOLabeler with three steps for AFP.

### 3.3 Component methods

We selected five typical, different sequence-based information for generating components. They are called *Naive* (GO term frequency), *BLAST-KNN* (B-K, *k*-nearest neighbor using BLAST results), *LR-3mer* (Logistic regression (LR) of the frequency of amino acid trigram), *LR-InterPro* (LR of InterPro features), and *LR-ProFET* (*LR* of *P*roFET features), which are all explained more below. These components from different information sources should be complement to each other.

#### 3.3.1 Naive: GO term frequency

For given *P*_*j*_, the score that *P*_*j*_ is with *G*_*i*_ can be computed simply by the frequency of *G*_*i*_ in *D*, as follows (Note that this method gives the same score for all P_j_):

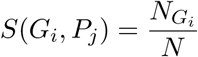

#### 3.3.2 BLAST-KNN (B-K): sequence alignment

It is reported that using the similarity score (bit-score) between similar proteins and the query slightly improves the performance of just using the sequence identity [25]. So for given *P*_*j*_, the score of BLAST-KNN *S(G_i_,P_j_)* is computed by first running BLAST to have the similarity score (bit-score) *B(Pj,p)* between *Pj* and protein p to identify a set *Hj* of similar proteins to *Pj* in D using a certain cut-off value (set at *e*-value of 0.001 in our experiments) against *B(P*_*j*_, *p*) for all *p*. Finally the score can be obtained as follows:

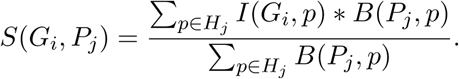

#### 3.3.3 LR-3mer: amino acid trigram

For each protein, we use the frequency of the types of three consecutive amino acids (amino acid trigram or 3mer) in D, which turns into a vector of 8,000 (= 20^3^) features. We then use this vector as an input of logistic regression classifier of each GO term.

#### 3.3.4 LR-InterPro: protein families, domains and motifs

InterPro [21] combines 14 different protein and domain family databases, including Pfam [28], CATH-Gene3D [27], CCD [20] and SUPERFAMILY [8], so covering a large number of protein families, domains and motifs in sequences. We run InterProScan^2^ over a sequence in D, resulting in a binary vector with 33,879 features, and then use this vector as an instance of training logistic regression classifier of each GO term.

#### 3.3.5 LR-ProFET: sequence features including biophysical properties

We run ProFET^3^ [23] over a sequence in *D* to extract features including elementary biophysical properties, etc., resulting in a vector of 1,170 features which is used as an input to train logistic regression classifier of each GO term.

### 3.4 GOLabeler

GOLabeler has three steps for functional annotation of a query protein (Fig. 1):

#### 3.4.1 Step 1: Generate candidates GO terms

For a query protein, we run five component methods to have predicted GO terms, and after choosing the top-*k* predicted GO terms from each component, merge them together as the candidate GO terms (we used *k*=30 in our experiments. See Section 4.3) Note that reducing *k* is to focus on the most relevant GO terms to query protein and also reduce the computational burden of the model.

#### 3.4.2 Step 2: Generate features for ranking GO terms

We then generate features of the query protein by using the scores (of each of the candidate GO terms) predicted by all five component methods, resulting in a 5-dimensional feature vector for each pair of a GO term and one query protein. Note that all score values are in between 0 and 1.

#### 3.4.3 Step 3: Rank GO terms by learning to rank (LTR)

Finally, we use LTR to rank all candidate GO terms of each query protein. Note that all proteins in the training data and their candidate GO terms are used for training the LTR model. By using LTR, we can effectively integrate multiple sequence-based evidence for AFP of *no-knowledge* Proteins.

### 3.5 Competing methods

In our experiments, we compare GOLabeler with five methods: three ensemble approaches with the same component outputs as GOLabeler: one vote, weighted voting (WV) and consensus (which have been often used in other AFP work, e.g. [30]), and two existing methods, BLAST [1] and GoFDR [10]. We note that GoFDR was a top performer of CAFA2.

#### 3.5.1 One vote

One vote selects the most confident prediction out of the five components.

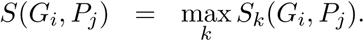

#### 3.5.2 Weighted voting (WV)

Weighted voting combines the predicted scores of component methods linearly by using weights over components as follows:

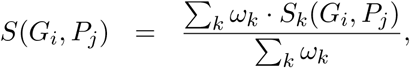

Where ω_*k*_ is the weight assigned to the *k*-th component. In our experiments, the weights are set in proportion to the area under precision-recall curve (AUPR) of each component. See Section 4.2 for AUPR more.

#### 3.5.3 Consensus

Consensus computes the score as follows:

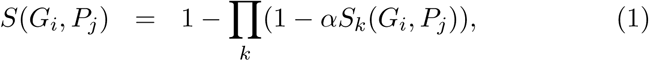

Where *a* ∈ [0, 1] is a constant to balance components by their importance. Here we use *α* = 1 that is most typical.

#### 3.5.4 **BLAST**

BLAST was used as a baseline method in both CAFA1 and CAFA2, and so we use this as a competing method. Similar to BLAST-KNN, given query protein *P*_*j*_, the similar proteins *H*_*j*_ in *D* to the query protein are obtained by using some cut-off value (again set at e-value of 0.001 in our experiments) against similarity score (bit-score) *B(P*_*j*_ *,p)* between *P*_*j*_ and protein *p* in *H*_*j*_, by which the score by BLAST can be obtained as follows:

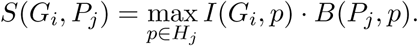

#### 3.5.5 GoFDR

Among the top performance methods in CAFA2, GoFDR is only the method having available source code^4^, and so we choose GoFDR as a competing method [10]. For a query protein, GoFDR run BLAST or PSI-BLAST to obtain multiple sequence alignment (MSA) over the query sequence and find functionally discriminating residues (FDR) of each GO term in the MSA which are used to generate a position-specific scoring matrix (PSSM). GoFDR uses the PSSM to compute the score between the query protein and a GO term.

## 4 Experiments

### 4.1 Data

Data collection approximately followed the corresponding part of CAFA1 [25] and CAFA2 [14]:

Protein sequences

We downloaded the FASTA-format files of all proteins from UniProt^5^ [29].

GO terms

We downloaded protein function annotation from SwissProt^6^ [3], GOA^7^ [12], and GO^8^ [2] in October 2016. Out of them we extracted all experimental annotations in: ‘EXP’, ‘IDA’, ‘IPI’, ‘IMP’, ‘IGI’, ‘IEP’, ‘TAS’, or ‘IC’, and then merged them to form a full annotation dataset (Note that SwissProt did not have annotation dates and so we downloaded data of SwissProt in January 2015 and January 2014 also).

We then generated the following four datasets, which are mainly separated by the time stamps that proteins are annotated.

Training: training for components

All data annotated in 2014 or before.

LTR1: training for LTR

Among the data experimentally annotated in 2015 and not before 2015, *no-knowledge* proteins.

LTR2: training for LTR

Among the data experimentally annotated in 2015 and not before 2015, *limited-knowledge* proteins. Note that the input of GOLabeler and other competing methods are sequences only, and so the input from LTR2 are also sequences only in our experiments.

Testing: testing for competing methods

All data experimentally annotated in 2016 (January to October of 2016, since we downloaded the data in October 2016) and not before 2016.

Note that this time-series way of separating training and testing data is the same as CAFA. Also we used the same target species as CAFA3, an ongoing challenge for AFP, in LTR1, LTR2 and Testing. Table 1 shows the number of proteins in the four datasets. We used Testing (or we call *benchmark)* as the test set to examine the performance of competing methods.

**Table 1:** Data statistics (# proteins) on species with at least ten proteins for one domain of GO for each of the four datasets.

| Species | Training |  |  | LTR1 |  |  | LTR2 |  |  | Testing |  |  |
| --- | --- | --- | --- | --- | --- | --- | --- | --- | --- | --- | --- | --- |
|  | MFO | BPO | CCO | MFO | BPO | CCO | MFO | BPO | CCO | MFO | BPO | CCO |
| HUMAN (*Homo sapiens*) | 9,087 | 11,019 | 15,977 | 150 | 216 | 718 | 261 | 454 | 161 | 42 | 64 | 67 |
| MOUSE (*Mus musculus*) | 5,697 | 9,262 | 8,770 | 169 | 339 | 237 | 140 | 137 | 175 | 100 | 235 | 146 |
| DROME (*Drosophila melanogaster*) | 4,646 | 1,0778 | 7,609 | 43 | 579 | 172 | 132 | 224 | 327 | 125 | 339 | 188 |
| ARATH (*Arabidopsis thaliana*) | 3,857 | 7,238 | 8,492 | 84 | 198 | 140 | 195 | 114 | 70 | 84 | 180 | 128 |
| DANRE (*Danio rerio*) | 2,173 | 8,374 | 1,500 | 76 | 722 | 159 | 108 | 56 | 50 | 66 | 380 | 27 |
| RAT (*Rattus norvegicus*) | 4,199 | 5,128 | 4,459 | 121 | 185 | 178 | 56 | 70 | 106 | 31 | 88 | 88 |
| DICDI (*Dictyostelium discoideum*) | 414 | 921 | 803 | 17 | 52 | 25 | 16 | 15 | 19 | 16 | 26 | 12 |
| All species (not only the above) | 45,543 | 77,170 | 71,388 | 724 | 2,387 | 1,679 | 1,081 | 1,206 | 956 | 497 | 1,340 | 770 |

### 4.2 Performance evaluation measures

We used three measures: AUPR (Area Under the Precision-Recall curve), F_max_ and S_min_ for evaluation of the predicted GO terms for each protein, i.e. a multi-label classification setting. AUPR as well as AUC (area under the receiver operator characteristics curve) are very general evaluation criteria for classification. AUPR punishes false positive more than AUC, resulting in being more frequently used when high costs are required for obtaining labels, such as experimental biology. F_max_ and S_min_ are less general but have been used in CAFA^9^. We explain F_max_ and S_min_ below (notation follows the Method section):

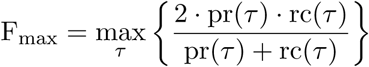

where pr(τ) and rc(τ) are so-called *precision* and *recall*, respectively, obtained at some cut-off value, τ, defined as follows:

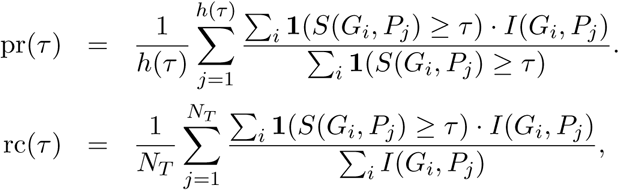

where *H(*τ) is the number of proteins with the score no smaller than τ for at least one GO term, and 1(-) is 1 if the input is true;otherwise zero.

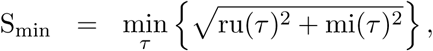

where ru(τ) and mi(τ) are two types of errors, called *remaining uncertainty* and *misinformation*, respectively, given as follows:

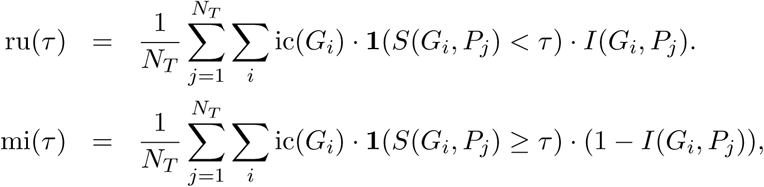

where ic(G,) is the *information content* of G*i*, defined as follows:

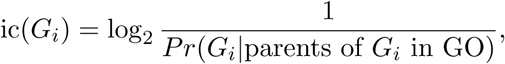

where P*r* (*G*_*i*_|parents of *G*_*i*_ in GO) is the conditional probability of *G*_*i*_ given its parents of the GO structure (see [5] for more details).

Simply AUPR, F_max_ and S_min_ will select methods which provide higher precision at top prediction, more balanced prediction between precision and recall, and more balanced prediction between two error types (and smaller errors for GO terms with higher information contents), respectively. Also note that practically we evaluated the top 100 GO terms predicted by competing methods for each GO domain (we used 100, since the number of GO terms per protein is clearly smaller than 100, and also simply the top GO terms are important).

### 4.3 Implementation and parameter settings

We processed the FASTA-format data by biopython^10^ and used sklearn^11^ for running logistic regression and xgboost [4] to run LTR.

#### 4.3.1 GOLabeler

BLAST-KNN

Ver. 2.3.0+ was used with default parameters for BLAST, except that blastdb was from all proteins in *D* and the number of iterations was one.

LTR

We used ‘rank:pairwise’ as the objective loss function in xgboost. Also the maximum depth of trees in MART (Multiple Additive Regression Trees) was set at 4, to avoid overfitting to the training data. We selected top 30 predictions from each component to be merged, since this number provided the most stable performance in five-fold cross validation over LTR training data (out of five values {10,20,30,40 and 50} tested).

#### 4.3.2 Competing methods

BLAST

We used the same setting as BLAST-KNN.

GoFDR

We ran GoFDR over all required data (i.e. all annotation without ‘IEA’ or ‘RCA’ evidence and some annotations with IEA evidence before 2016 from GOA [12]).

### 4.4 Results

We resampled the test dataset with replacement 100 times (bootstrap with replacement) to make the experiment reliable. In addition to the three performance evaluation measures, we used paired *t*-test to statistically evaluate the performance difference between the best performance method (in boldface in tables) and all other methods, where the result was considered significant if *p*-value was smaller than 0.05. Then in tables, the best performance value is underlined if the value is statistically significant (see the supplementary materials for the detailed *p*-values).

#### 4.4.1 Comparison with component methods

We first compare the five component methods, the results being shown in Table 2, where out of the five methods, the best values in each column are in italics with asterisks. Out of the nine (= three evaluation criteria times three domains) comparison settings, LR-InterPro achieved the best five times, being followed by BLAST-KNN which achieved the best three times, implying that LR-InterPro and BLAST-KNN are the two best component methods. The other three methods are less accurate, while their high performance could be found in some specific case: LR-3mer achieved the highest Fmax in CCO. We then examined the performance of GOLabeler trained by LTR1, the results being shown in the same table, where GOLabeler with only two components was also checked as well as GOLabeler (with all five components), which are called *GOLabeler (B+I)* and *GOLabeler (All)*, respectively. The table shows that GOLabeler (All) outperformed all competing methods in eight out of nine evaluation settings. For example, GOLabeler (All) achieved the highest F_max_ of 0.580 in MFO, followed by GOLabeler (B+I) of 0.578, BLAST-KNN of 0.573 and LR-InterPro of 0.556. This result indicates the advantage of incorporating all component methods in GOLabeler, compared with using only a smaller number of components. Another finding was that among MFO, BPO and CCO, BPO is the hardest task in AFP (this is consistent with the results of CAFA). For example, the best AUPR of GOLabeler (All) was 0.546 and 0.700 for MFO and CCO, respectively, while it was only 0.225 for BPO, implying that sequences are the limited informative for BPO in AFP. Hereafter, we show the result by GOLabeler (All) as that by GOLabeler.

**Table 2:**
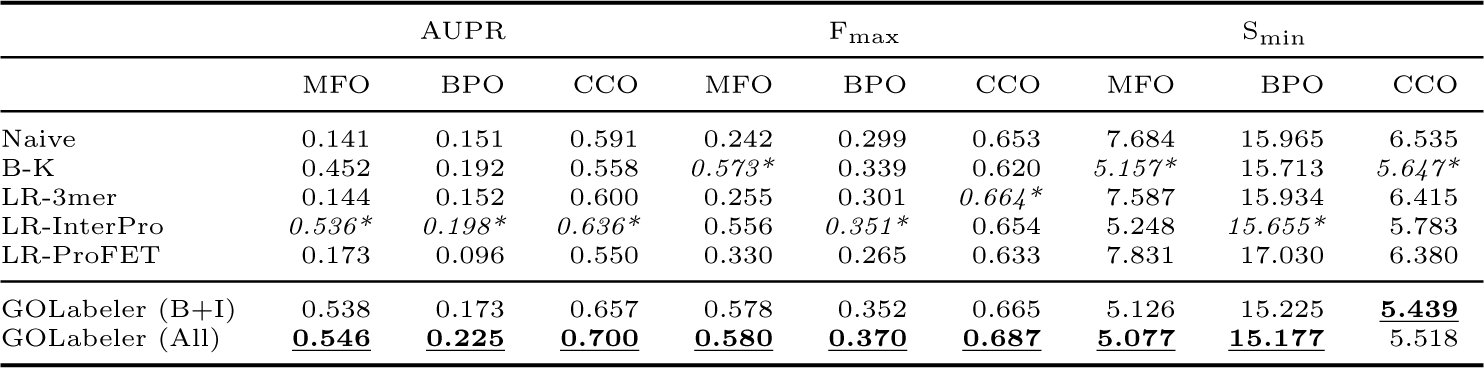
Performance comparison of GOLabeler with component methods. B-K: BLAST-KNN. GOLabeler (B+I): only BLAST-KNN and LR-InterPro were used for components in GOLabeler. GOLabeler (All): all five components are used.

|  | AUPR |  |  | F<sub>max</sub> |  |  | S<sub>min</sub> |  |  |
| --- | --- | --- | --- | --- | --- | --- | --- | --- | --- |
|  | MFO | BPO | CCO | MFO | BPO | CCO | MFO | BPO | CCO |
| Naive | 0.141 | 0.151 | 0.591 | 0.242 | 0.299 | 0.653 | 7.684 | 15.965 | 6.535 |
| B-K | 0.452 | 0.192 | 0.558 | *0.573\** | 0.339 | 0.620 | *5.157\** | 15.713 | *5.647\** |
| LR-3mer | 0.144 | 0.152 | 0.600 | 0.255 | 0.301 | *0.664\** | 7.587 | 15.934 | 6.415 |
| LR-InterPro | *0.536\** | *0.198\** | *0.636\** | 0.556 | *0.351\** | 0.654 | 5.248 | *15.655\** | 5.783 |
| LR-ProFET | 0.173 | 0.096 | 0.550 | 0.330 | 0.265 | 0.633 | 7.831 | 17.030 | 6.380 |
| GOLabeler (B+I) | 0.538 | 0.173 | 0.657 | 0.578 | 0.352 | 0.665 | 5.126 | 15.225 | **5.439** |
| GOLabeler (All) | **0.546** | **0.225** | **0.700** | **0.580** | **0.370** | **0.687** | **5.077** | **15.177** | 5.518 |

#### 4.4.2 Comparison with other ensemble techniques and also performance improvement by making training size larger

We examined the performance of recent ensemble techniques with the same five component methods and also the performance improvement by increasing the training data for ensemble learning, i.e. weighted voting and GOLabeler. Table 3 shows the result summary under nine experimental settings. When we used LTR1 only, GOLabeler achieved the best performance among the competing methods. Weighted voting was the next best, while the performance of one vote was lowest in all nine settings, implying that choosing one component would not work well. This result is inconsistent with that of [30] but might be reasonable because of the large difference of datasets. In fact, [30] assumes that functional annotation within the same ortholog group (OG) should be shared and transferable, focusing on only 4,145 GO terms. On the other hand, we followed the experimental setting of CAFA, generating a huge number of GO terms. Thus, in our case a single classifier would not make better prediction than ensembles, by which our results would make sense.

**Table 3:**
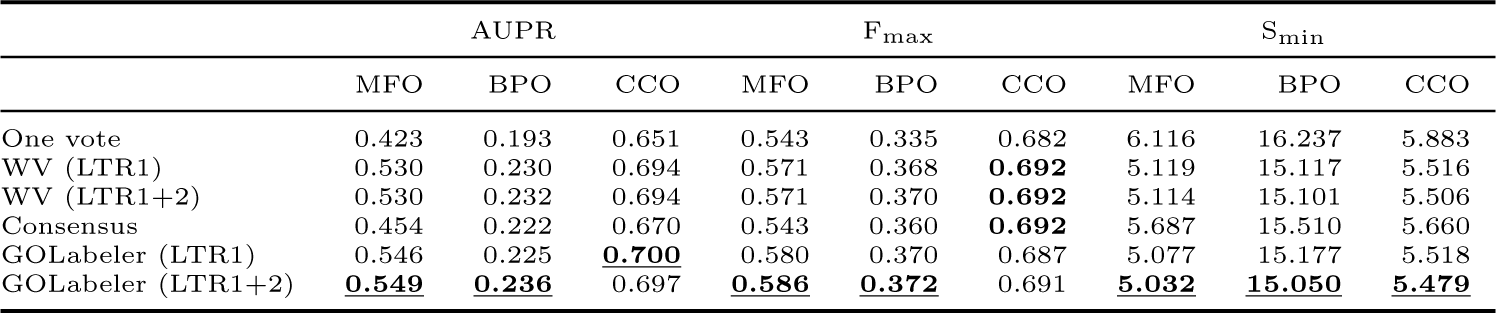
Performance comparison of GOLabeler with ensemble techniques and also improvement by adding LTR2 to LTR1. WV: Weighted voting.

|  | AUPR |  |  | F<sub>max</sub> |  |  | S<sub>min</sub> |  |  |
| --- | --- | --- | --- | --- | --- | --- | --- | --- | --- |
|  | MFO | BPO | CCO | MFO | BPO | CCO | MFO | BPO | CCO |
| One vote | 0.423 | 0.193 | 0.651 | 0.543 | 0.335 | 0.682 | 6.116 | 16.237 | 5.883 |
| WV (LTR1) | 0.530 | 0.230 | 0.694 | 0.571 | 0.368 | **0.692** | 5.119 | 15.117 | 5.516 |
| WV (LTR1+2) | 0.530 | 0.232 | 0.694 | 0.571 | 0.370 | **0.692** | 5.114 | 15.101 | 5.506 |
| Consensus | 0.454 | 0.222 | 0.670 | 0.543 | 0.360 | **0.692** | 5.687 | 15.510 | 5.660 |
| GOLabeler (LTR1) | 0.546 | 0.225 | **0.700** | 0.580 | 0.370 | 0.687 | 5.077 | 15.177 | 5.518 |
| GOLabeler (LTR1+2) | **0.549** | **0.236** | 0.697 | **0.586** | **0.372** | 0.691 | **5.032** | **15.050** | **5.479** |

**Table 4:**
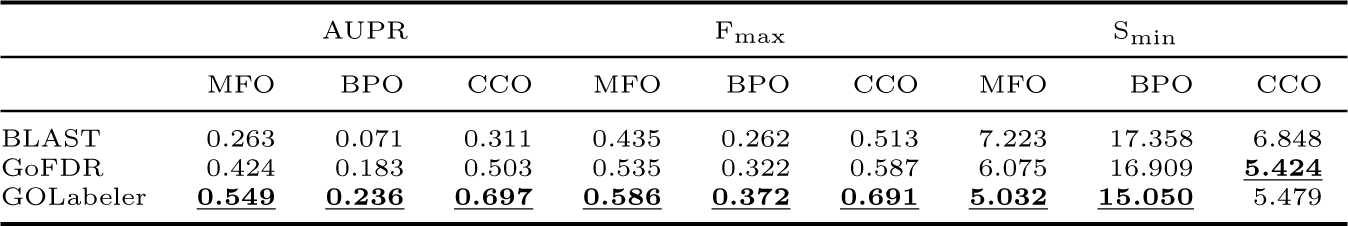
Performance comparison of GOLabeler with BLAST and GoFDR.

By adding LTR2, the performance of GOLabeler was more pronounced, achieving the best in all nine settings, except only one, while the performance of weighted voting was increased in only five out of all nine cases. This means that GOLabeler can take the advantage of using a larger size of data more effectively than weighted voting. Also this implies that the performance of GOLabeler can be rather easily improved further by increasing the annotation data in the future. Hereafter, we show the results of GOLabeler and weighted voting with all training data for LTR (both LTR1 and LTR2) as those by GOLabeler and weighted voting, respectively.

#### 4.4.3 Comparison with BLAST and GoFDR

Table 5 shows the performance of BLAST and GoFDR. GOLabeler outperformed both BLAST and GoFDR in all experimental settings, except only one, being statistically significant. GoFDR is a method that achieved the top performance in CAFA2, demonstrating the high performance of GOLabeler, even compared with high performers in CAFA.

**Table 5:**
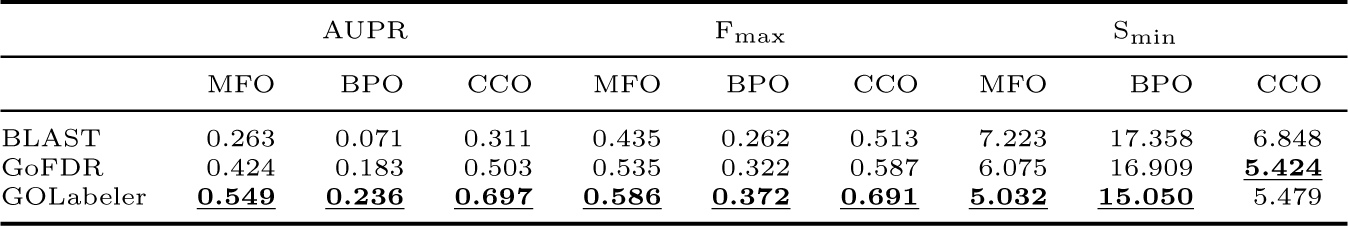
Performance comparison of GOLabeler with BLAST and GoFDR.

#### 4.4.4 Comparison with BLAST and GoFDR

Table 5 shows the performance of BLAST and GoFDR. GOLabeler outperformed both BLAST and GoFDR in all experimental settings, except only one, being statistically significant. GoFDR is a method that achieved the top performance in CAFA2, demonstrating the high performance of GOLabeler, even compared with high performers in

#### 4.4.5 Performance over different species

We checked the performance variation depending on different species, particularly focusing on two species with the largest number of proteins (over 100 proteins in all three GO domains), i.e. DROME and MOUSE, for which results are summarized in Table. 6 and 7, respectively. The two tables show that GOLabeler achieved the best performance in all nine settings for DROME, except one case, while GOLabeler was the best method for MOUSE but the best cases remained four out of all nine settings. We can see species-wise slight performance variation, although performance advantage of GOLabeler was clear.

**Table 6:**
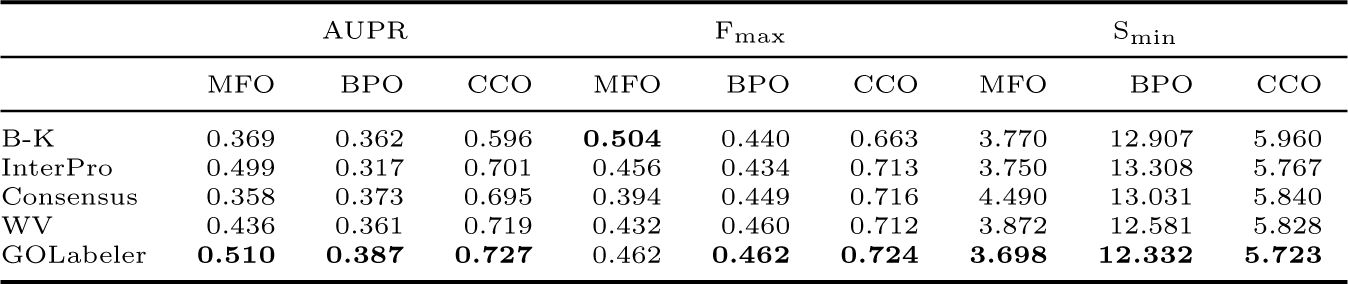
Performance comparison, focusing on DROME. B-K: BLAST-KNN. WV: Weighted voting.

**Table 7:**
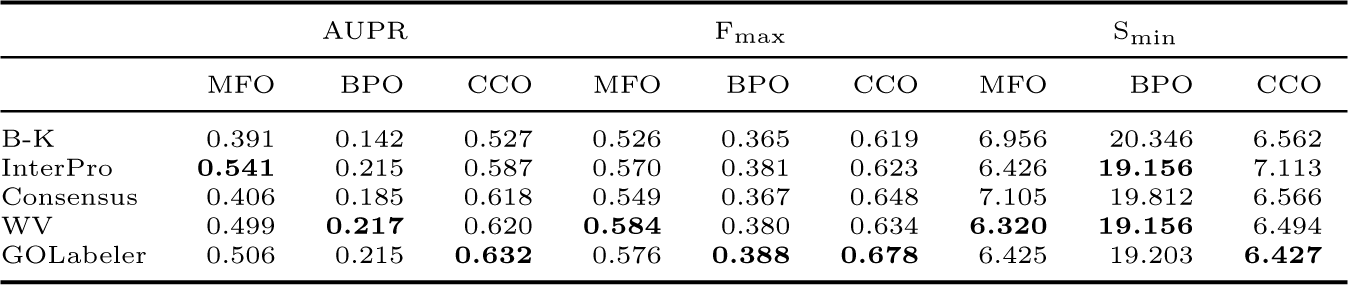
Performance comparison, focusing on MOUSE. B-K: BLAST-KNN. WV: Weighted voting

|  | AUPR |  |  | F<sub>max</sub> |  |  | S<sub>min</sub> |  |  |
| --- | --- | --- | --- | --- | --- | --- | --- | --- | --- |
|  | MFO | BPO | CCO | MFO | BPO | CCO | MFO | BPO | CCO |
| B-K | 0.391 | 0.142 | 0.527 | 0.526 | 0.365 | 0.619 | 6.956 | 20.346 | 6.562 |
| InterPro | **0.541** | 0.215 | 0.587 | 0.570 | 0.381 | 0.623 | 6.426 | **19.156** | 7.113 |
| Consensus | 0.406 | 0.185 | 0.618 | 0.549 | 0.367 | 0.648 | 7.105 | 19.812 | 6.566 |
| WV | 0.499 | **0.217** | 0.620 | **0.584** | 0.380 | 0.634 | **6.320** | **19.156** | 6.494 |
| GOLabeler | 0.506 | 0.215 | **0.632** | 0.576 | **0.388** | **0.678** | 6.425 | 19.203 | **6.427** |

#### 4.4.6 Performance over different GO terms

We have evaluated the performance of competing methods regarding if GO terms are predicted correctly for each protein, which is a protein-centric manner. We now evaluated the competing methods in terms of if true proteins are correctly predicted for each GO-term, which is a term-centric manner, or more generally binary classifiication, by using AUPR. We focused on GO terms which appear no less than 10 times in the test data. Note that GOLabeler was trained to optimize the ranking of GO terms per protein, but even under this case, the result shows that GOLabeler achieved the highest average AUPR of 0.659 in MFO, being followed by weighted voting of 0.646 and consensus of 0.638. Table 8 shows the top five GO terms in MFO on the difference in AUPR of GOLabeler from that of the next best method (the difference was all larger than 4% in AUPR).

**Table 8:** Top five GO IDs in MFO by GOLabeler outperformed all other methods

| ID | description | AUPR |
| --- | --- | --- |
| GO:0052689 | carboxylic ester hydrolase activity | 0.697 |
| GO:0022891 | substrate-specific transmembrane transporter activity | 0.832 |
| GO:0043565 | sequence-specific DNA binding | 0.614 |
| GO:0022892 | substrate-specific transporter activity | 0.775 |
| GO:0043169 | cation binding | 0.606 |

#### 4.4.7 Case study

Finally we show a typical example of the results obtained by GOLabeler and other competing methods, to illustrate the real effect of the performance difference on annotating GO to unknown proteins. Table 9 shows the list of predicted GO terms^12^ of MFO for a protein, Wor4p (Uniprot Symbol: Q5ADX8), which is, in ground-truth, associated with 12 GO terms in MFO. Also Fig. 2 shows the directed acyclic graph of the 12 GO terms associated with Q5ADX8. First note that there are no homologous proteins of Q5ADX8 (set the cut-off of *e*-value at 0.001), by which BLAST-KNN was unable to predict any GO terms. Naive predicted three GO terms, out of which only one very generally term, GO:0005488 (Binding), was correct. LR-InterPro, LR-3mer, LR-ProFET could make more correct annotations particularly more specific terms, such as GO:0003677 (DNA Binding). The prediction by weighted voting was also at the same level of specificity as the three component methods^13^. In fact, the predicted GO terms by LR-InterPro and weighted voting were all correct, but the number of the predicted GO terms was only four and five, respectively. On the other hand, GOLabeler predicted nine terms, out of which eight were correct. More importantly, as shown in Fig. 2, the prediction by GOLabeler was most specific. For example, even GO:0044212 (transcription regulatory region DNA binding), which is next to the end node in Fig. 2, was correctly annotated. From this result, we can see that the performance advantage of GOLabeler results in sizeable differences in quality of real function annotation.

**Table 9:** Predicted GO terms of Q5ADX8 in MFO by GOLabeler and competing methods. Correctly predicted GO Terms are in bold face. WV: Weighted voting.

| Method | Predicted GO terms |
| --- | --- |
| Naive | **GO:0005488**, GO:0003824, GO:0005515 |
| BLAST-KNN |  |
| LR-3mer | **GO:0005488**, GO:0005515, GO:0003824, **GO:0097159**, **GO:1901363**, **GO:0003676** |
| LR-InterPro | **GO:0005488**, **GO:0003677**, **GO:1901363**, **GO:0097159**, **GO:0003676** |
| LR-ProFET | **GO:0003676**, **GO:0097159**, **GO:1901363**, **GO:0005488**, GO:0003723, **GO:0003677**, GO:0001010, GO:0005515 |
| WV | **GO:0005488**, **GO:0097159**, **GO:1901363**, **GO:0003676** |
| GOLabeler | **GO:0005488**, **GO:0003676**, **GO:0097159**, **GO:1901363**, **GO:0003677**, GO:0005515, **GO:0044212**, **GO:0000975**, **GO:0001067** |
| Ground truth | GO:0005488, GO:0097159, GO:1901363, GO:0003676, GO:0001067, GO:0003677, GO:0000975, GO:0003690, GO:0043565, GO:0044212, GO:1990837, GO:0000976 |

**Figure 2:**
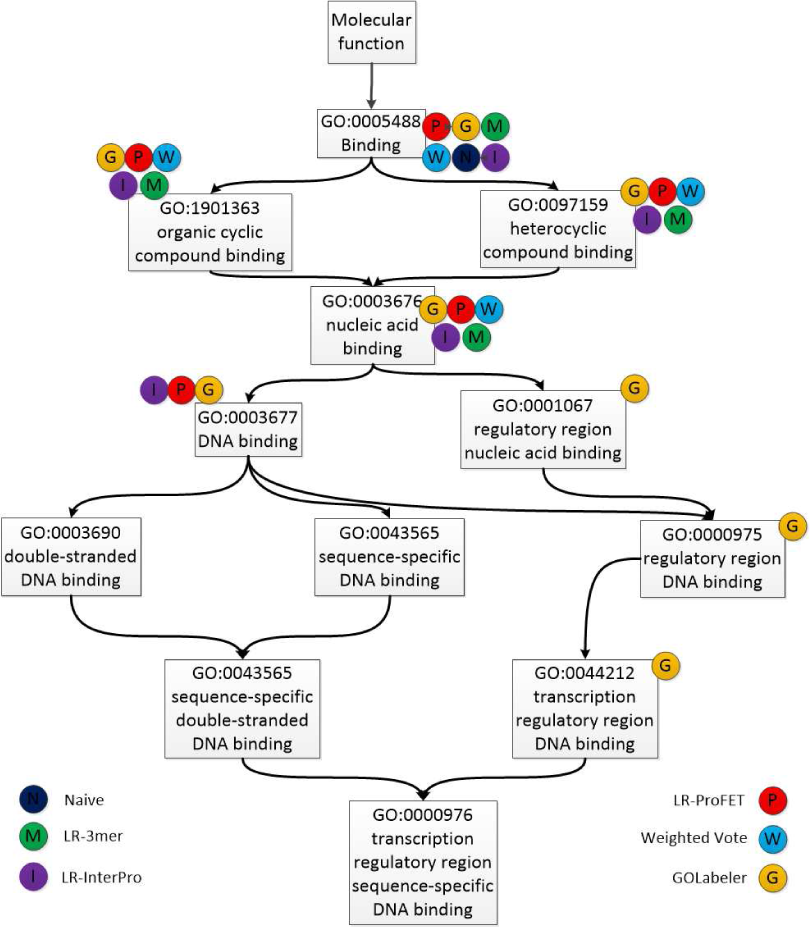
Predicted GO terms of Q5ADX8 in DAG of MFO by different methods.

## 5 Conclusion

Sequence-based large-scale automated function prediction (SAFP) for proteins, particularly *difficult* proteins, is an important and challenging problem. Many factors have made this problem difficult to solve. First of all, in this setting, experimental data except sequences cannot be used, while difficult proteins have very low sequence homology to other sequences in training data, by which homology-based approaches do not work. Also GO protein functions themselves have mainly two issues: 1) in a directed acyclic graph (DAG) of GO, a protein with some GO terms must be annotated by all GO terms of ancestor nodes. 2) So the GO structure resulting in a huge bias in the number of GO terms per protein, making AFP a hard multi-label classification problem. GOLabeler addresses these challenges by cutting-edge techniques, particularly those in machine learning. 1) We use not only sequence homology, e.g. sequence alignment, but also a variety of sequence features, such as n-gram, domains, motifs, biophysical properties, etc., as features to train multiple classifiers, such as *k*-nearest neighbors (local information) and logistic regression (rather global information). 2) We use all GO terms, which are generated by considering the DAG structure of GO, as class labels of classifiers. 3) Finally the well-trained classifiers are integrated as components by a cutting-edge machine learning technique, learning to rank (LTR), which can rank the candidate GO terms in terms of the relevance to each protein.

We have thoroughly examined the performance of GOLabeler extensively, showing the clear advantage in predictive performance over state-of-the-art techniques in AFP and ensemble approaches. Again using diverse information from sequences is very useful for AFP. GOLabeler currently integrates five classifiers as components which are from GO term frequency, sequence homology, trigram, motifs and biophysical properties, and so on. The framework of GOLabeler or LTR is flexible and allows to incorporate any classifiers we can use, implying that further performance improvement would be possible by adding more information from sequences. Also this could be done for not only SAFP but also more general AFP. This would be interesting future work. From our experimental results, we have seen the noteworthy performance improvement of GOLabeler for difficult proteins. Also we saw the slight performance difference depending on species. Possible future work would be to develop GOLabeler to improve the performance of AFP for specific species, especially those with very few proteins, for which current AFP methods do not necessarily work sufficiently.

## Footnotes

1 Also currently (2016–2017), CAFA3 is on-going.

2 http://www.ebi.ac.uk/interpro/interproscan.html

3 https://github.com/ddofer/ProFET

4 http://gofdr.tianlab.cn

5 http://www.uniprot.org/downloads

6 http://www.uniprot.org/downloads

7 http://www.ebi.ac.uk/GOA

8 http://geneontology.org/page/download-annotations

9 Evaluation criteria have been actively discussed, e.g. [13].

10 http://biopython.org/

11 http://scikit-learn.org/stable/index.html

12 The GO terms by each method are determined by its own cut-off value to achieve the best value of F_max_.

13 The poor performance of weighted voting might be caused by BLAST-KNN, which was unable to predict any GO term.

